# Attacked from above and below, new observations of cooperative and solitary predators on roosting cave bats

**DOI:** 10.1101/550582

**Authors:** Krizler Tanalgo, Dave Waldien, Norma Monfort, Alice Hughes

## Abstract

Predation of bats in their roosts has previously only been attributed to a limited number of species such as various raptors, owls, and snakes. However, in situations where due to over-crowding and limited roost space, some individuals may be forced to roost in suboptimal conditions, such as around the entrances of caves and may thus be vulnerable to predation by species which would normally be unlikely to predate bats whilst roosting inside caves. Here, we describe the first documented cooperative hunting of the Large-billed Crow, *Corvus macrorhynchos* Wagler, 1827 (Passeriformes: Corvidae) and opportunistic predation by the Yellow-headed water monitor, *Varanus cumingi* Martin, 1839 (Squamata: Varanidae) in the world’s largest colony of Geoffroy’s Rousette, *Rousettus amplexicaudatus* (É. Geoffroy Saint-Hilaire, 1810) (Chiroptera: Pteropodidae) in the Island of Samal, Mindanao, Philippines.

## 1. Introduction

Animal feeding behaviour and strategy i.e., to locate, acquire, and consume food resources shaped their characteristics, community interactions, and the development of their complex behaviour to efficiently obtain food and survive (Roberts 1942, McFarland 1981). While, the availability of resources and predation are among the most important biotic drivers that shape interactions across wildlife communities (Talbot 1978, Hiltunen and Laakso 2013; Glen and Dickman 2014). Within an ecosystem, when the predator populations decline or are absent, the prey population may dramatically increase (Gese and Knowlton 2001, Ritchie at al. 2013). This release of a species numbers may have negative implications especially when the prey is an invasive and out-compete native species for space and other resources (Ritchie at al. 2013) therefore, predator plays a crucial role to balance and shape the prey population. Predation is a complex behaviour and can involve many different approaches, these include (1) ‘cooperation’ when hunting occurs in a pair or group, (2) ‘cheating’ when hunters spot prey but only join at the end of a hunt (3) ‘scavenging’ when hunters do not hunt but join whenever it has a chance to take food or when predators consume carrion (DeVault at al. 2003), and (4) ‘solitary’ where predator hunts alone (Packer and Ruttan, 1988).

Until a recent global review, the extent of bird predation on bats was not widely known (Mikula at al. 2016). Diurnal birds particularly raptors prey on at least 124 species (11 families) of bats (Mikula at al. 2016). Hunting and predation by birds frequently occur near caves and roosting areas where bats are abundant and predators can hunt bats at emergence and during foraging (Laycock 1982, Lee and Kuo, 2001, Stimpson 2009). Inside caves and underground ecosystems, reptiles particularly snakes are widely known predators for cave-dwelling bats while varanids (monitor lizards) have not previously been reported. In the Asian tropics, few bat predators have been documented and there may be more interactions unreported (Lima and O’Keefe 2013, Mikula at al. 2016). In the Philippines, large Raptors (Family: Accipitridae) and pythons have been noted to prey on tree roosting flying fox *Acerodon jubatus* (van Weerd at al. 2003) nonetheless this is the first clearly documented and described observation of crow predation on a cave-dwelling bat (Tanalgo and Hughes 2018). In this paper, we describe and explain the hunting behaviour of two main predators of the cave-dwelling bat *Rousettus amplexicaudatus* in Monfort bat cave, Mindanao, Philippines: the Large-billed Crow (*Corvus macrorhynchos*) and Yellow-headed water monitor (*Varanus cumingi*).

## 2. Observations

We conducted a field observation on the Island of Samal located within the Gulf of Davao in the southern part of the Philippine Archipelago at the world’s largest colony of the Geoffroy’s Rousette (*Rousettus amplexicaudatus,* Chiroptera: Pteropodidae) (estimated population at present 2 million) (Carpenter at al. 2014) (Figure 1A) in the Monfort Bat Cave (7°09’53”N, 125°41’31”E). The rousette bat is a medium-sized cave-dwelling fruit bat (Figure 1B) native to Southeast Asia and widely distributed throughout the Philippines (Csorba et al. 2008; Heaney et al. 2010). Monfort Bat Cave is located in the private Monfort Bat Sanctuary and was opened for external ecotourism in 2005. Scientific research is allowed and several studies have been made before. Monfort Bat Cave is a relatively small cave with an approximate size of 150-metre length with an average 3 metres high and 5 metres wide. The cave system can be accessed through its horizontal entrance (opening #1) and four vertical openings (Carpenter at al. 2014).

**Figure 1:**
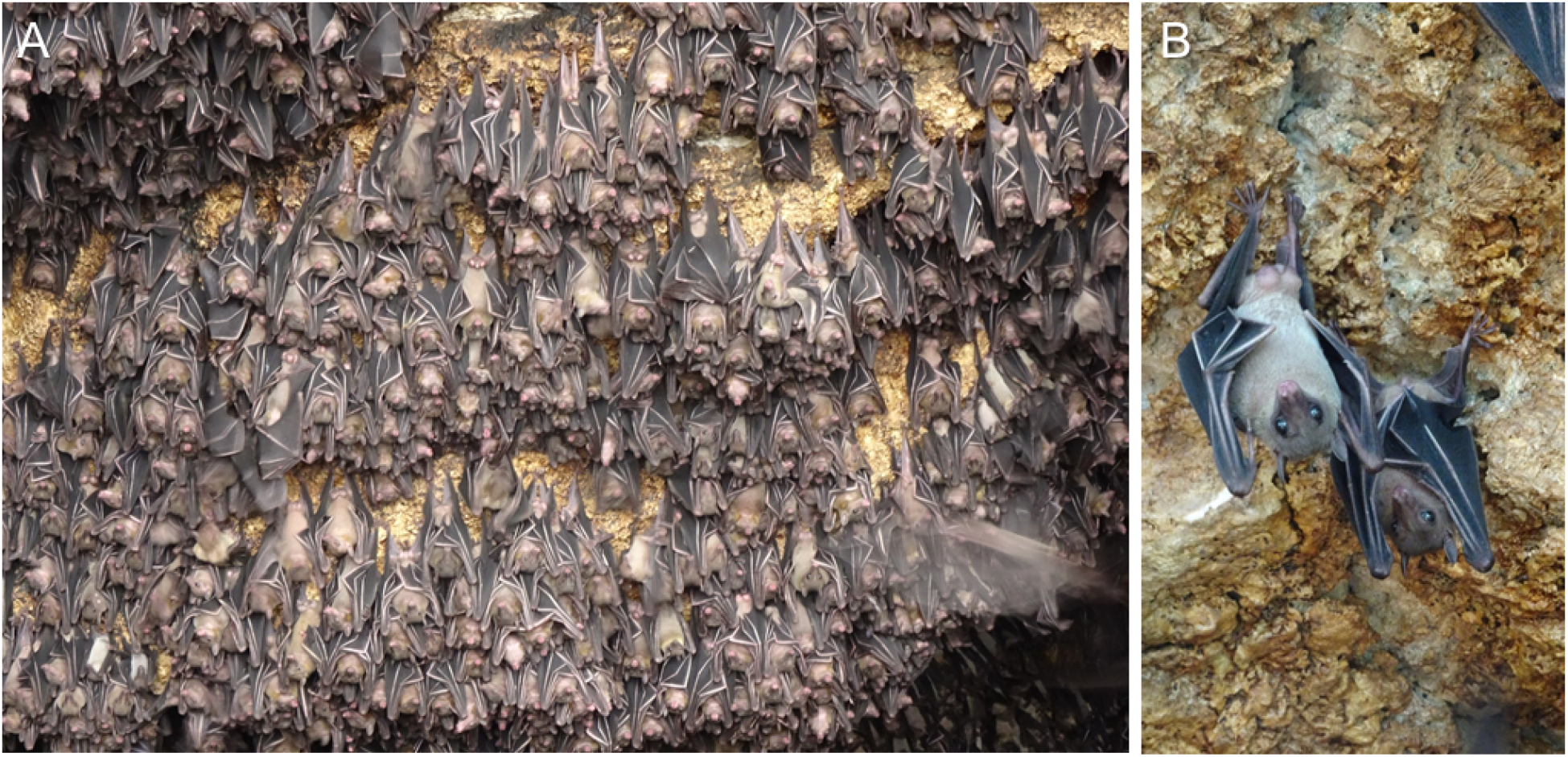
A. Large colony of R. amplexicaudatus in Monfort Cave in Samal Island, Philippines, B. Close-up image of *R.amplexicaudatus*

### 2.1. Cooperative predation of large-billed crow (Corvus macrorhynchos)

In Monfort cave, crows have been initially known to predate on bats since 2006 but their behaviour is not recorded and described in-depth. In late 2018 (September to December) we recovered decomposing carcasses of *R. amplexicaudatus* approximately 5-10 metres away from the entrances of five chambers of the cave. Further, we repeatedly observed crows perching on the trees surrounding the cave and flying into the cave preying upon the bats. In the succeeding months, we documented and described the patterns of their hunting behaviour. We observed crow activity 50 metres away from the cave opening and reinforced our observation by strategically installing a single wireless microcamera (TT microcamera™, P.R. China coupled with TT Android-based application support) at the first and third cave openings starting from 0700 H to 1700 H. Over 60 observational hours were made in the study site.

We observed 10 to 15 crows (*Corvus macrorhyncus*) flocking on trees 10 metres away from the caves. Two groups then form two separate groups: the first group is the standby-group (designated as “group A”) with ~9 to 10 individuals in flock, perching on tree branches around the caves while the second is the hunting group (designated as “group B”) with smaller number in flock with ~2-7 individuals. The on-standby group while remaining alert for potential disturbance and threats (e.g., human presence), the two crow murders occasionally produce loud calls apparently communicating with the groups. The hunting group (group B) positions (surveillance phase) on the fence or vines surrounding the cave openings locating the population to strategise the attack. This phase is the longest that occasionally lasted up to an estimated 10-15 minutes. After this phase, when crows in group B already identified its target group, it will separate to 2 subgroups to begin the hunt. The third phase (hunting) is described as “divide and conquer”. It begins when the first subgroup (dividers) of crow hunters flew swiftly descending towards the cave entrance, hovering, and mobbing the roosting bats to distract the colony population in the uppermost to mid-part part of the cave openings causing the individuals to dilute, which then other subgroups of hunters (conquerors) take advantage and easily capturing the prey bats. The hunting group then forcefully seize the bats using their strong beaks and ascend from the cave. This phase takes a maximum time of 8.78 minutes with an average of 2.20 minutes (*n*=26 hunting bouts). The hunting phase is not always successful in the first attempt and takes another second to third attempts for the same hunters to successfully capture a prey. In the last phase (re-grouping), group A joins the group B to take their share by assisting the latter group to rip apart the still active prey. Typically, during each of the 2-3 attempts hunting crows could successfully kill single bat individual (graphically described in Figure 2). In some observations, when a bat escapes seizure from group B and wings outside the cave, group A that is on stand-by tails and attack the bat and rip it apart with the member of the same group.

**Figure 2:**
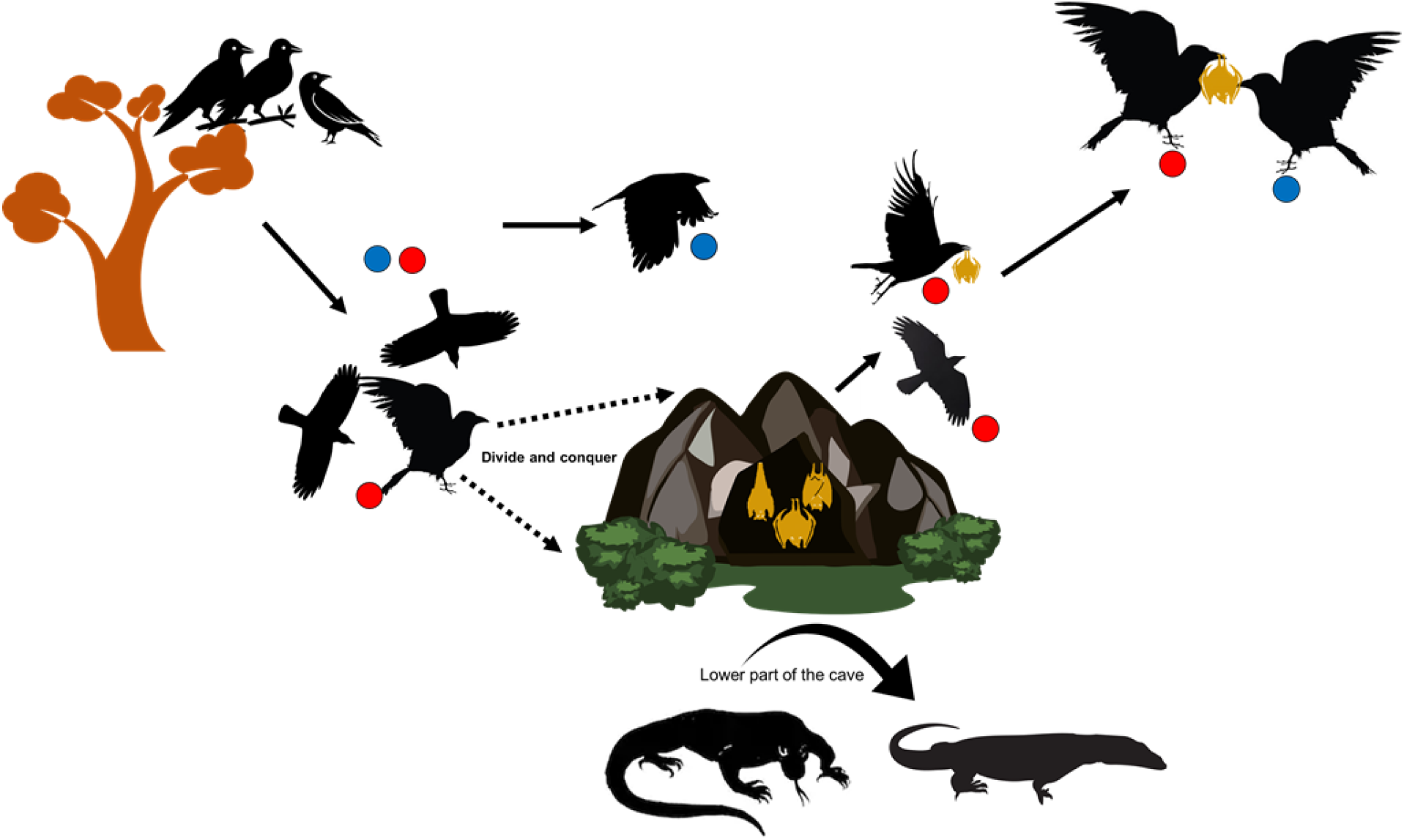
A diagram illustrating the cooperative hunting in crows (*C. macrorrhyncos*) (A-E) and solitary hunting of water monitor *V. cumingi* on cave-dwelling bat Rousettus amplexicaudatus in Samal Island, Philippines.

Crow hunting starts to be active as soon as the sunrise from 0600 H and recurs throughout the day and each hunting phases are variable but the relative length is consistent until 1600 H to 1700 H before the bat emergence and began to dark. The cave is intended for ecotourism activities and research, therefore, human presence is highly visible in the vicinity and this challenges the precision of the measurement of hunting recurrence or length per day as crows may disappear and pause bat hunting when tourists or the observers are noticeably present (e.g., numerous groups of tourist are present and noisy). Noticeably, days with a larger number of tourists the incidence of crow hunting bouts is lower (between 2 to 3 times per hour) compared to 6-8 times per hour on days with a lower number of visiting tourists. Additionally, because the exact population of crows we observed in the study is not certain we cannot determine if the same individuals perform the same role. The hunting success rate of crows are frequently successful but the distribution of food (e.g., amount of meat) varies and mostly favours the group B (hunters).

### 2.2. Solitary hunting of Yellow-headed water monitor (Varanus cumingi)

In addition to cooperative hunting by large-billed crows, we also documented solitary opportunistic predation by *V. cumingi* on cave bat *R. amplexicaudatus.* Unlike other Philippine endemic varanids (e.g., *Varanus olivaceus, V. mabitang*, and *V. bitatawa*) that are frugivorous (Koch at al. 2010, Law at al. 2018), *V. cumingi* is chiefly carnivorous and common in mangroves, fishponds, and tropical forests (Losos and Greene 1988, Koch et al. 2007, Sy et al. 2009). According to historical accounts, *V. cumingi* are resident cave dweller and chiefly remain inside, rarely venturing out where they are vulnerable to hunting (Tata Albeti pers. comm.). We observed two large *V. cumingi* preying solitarily on roosting bats in the lower part of the cave (cave opening #3 and #4) where less active (adult, sleeping, or ill) bats are found. The predation behaviour of *V. cumingi* is solitary, swift, less sophisticated but more efficient compared to cooperative hunting by crows. In our observation, a single varanid starts the hunting with surveillance, where a single individual water monitor lurks from boulders below the cave, moving its head left and right, and slowly creeping towards its identified colonies to prey (duration: ~2 to 3 minutes). When the monitor finds its potential prey, it immediately attacks the prey and swallows the whole single individual (duration: ~5-8 minutes). After consuming its prey, it slowly creeps back and rests under the boulders. Based on cave morphology, patterns of bats roosting in the cave, and likely behaviour of *V. cumingi*, we believe the lizard preys upon bats roosting on the lower cave walls as well as on and underneath the rock breakdown in a cave. They are likely more successful preying upon vulnerable bats that are injured, sick, older, or younger. There are instances of smaller varanids attempting to capture bats in the upper part walls but these frequently fail. Conspecific encounters were not observed during the course of our work.

## 3. Discussion

This is the first description of the unusual hunting behaviour by large-billed crow *C. macrorrhyncos* and *V. cumingi* on a cave-dwelling bat species, *R. amplexicauda*tus. The *C. macrorhyncus* is a monogamous cosmopolitan species widely distributed throughout the Asian region (Marzluff and Angell 2005, Tanalgo et al. in press). These species are known omnivorous but they tend to be carnivorous when animal prey are abundantly available (Goodwin 1982, Richardson and Verbeek 1997, Tanalgo at al. 2015). Large-billed crows was documented to prey on domestic cats (*Felis catus*), murids, rock doves (*Columba livia*), Grey Starlings (*Sturnus cineraceus*), and Tree Sparrows (*Passer montanus*) (Kurosawa at al. 2003). In Southeast Asia, forest bat *Cynopterus sphinx* are preyed by *C. macrorhyncus* and its commonly co-occurring species *C. enca* is known to prey on *Tadarida plicata* (as accounted by Mikula at al. 2016). Also, *C. brachyrhynchos* was previously recorded in temperate region preying on cave-dwelling bats *Myotis lucifugus* but only during flight and emergence (Lefevre 2005, Henandez at al. 2007, Flaspohler 2017). Although past evidence suggest that *C. macrorrhyncos* are proficient of the sophisticated cooperative and coordinated hunting the present observed hunting behaviour is peculiar and never been observed before, particularly for non-trivial bat prey. Cooperative hunting is an opportunistic or simultaneous hunting behaviour documented in few vertebrate groups (Bernard 1988, Packer and Ruttan 1988, Dinets 2017). This occurs when two or more individuals or groups of conspecifics cooperate to hunt larger or more aggressive prey to increase hunting success while reducing energy expenditure (Hector 1986, Bernarz 1988, Bowman at al. 2003). This strategy is noticeably identified on apparently observed coordination of the group or flocks, for example, each member attacks the target prey sequentially, which the hunters allow others groups to take advantage situation and benefit (Dinet 2015). Cuban boas (*Chilabothrus angulifer*), for example, was documented to block cave bats in cave opening to increase hunting efficacy giving an advantage to other groups to hunt (Dinets at al. 2017). In our present observation, the division of tasks among crow murders were apparently observed. The hunting groups, for instance, mob and disturb prey groups to become more vulnerable and easier for other participating hunters to seize and, carry the prey for other groups to assist in ripping apart for food sharing. Many studies have observed and suggests the complex intelligence and social behaviour of corvids (Emery and Clayton 2004) and cooperative behaviour is widely described in many species of in the wild (e.g., Bowman 2003, Yosef and Yosef 2010). Our present observation similar in pattern in hunting behaviour of *C. ruffi-collins* on Egyptian mastigure *Uromastyx aegypticus*, where crows are aware of their roles, divide into groups to perform coordinated and cooperative hunting of the prey (see Yosef and Yosef 2009).

However, despite hunters share resources, cooperative hunting strategy does not always assure higher food intake (e.g., Mech and Boitani 2003 demonstrated on grey wolf) or if the prey is smaller, only those aggressive and efficient hunters can take advantage of higher food share and intake (Packer and Ruttan 1988). Our present observation showed that food intake mostly favours the hunting groups (group B) compared to those in standby groups (group A). On the other hand, *Varanus cumingi* hunting strategy is less complex compared to the hunting strategy of crows. It is abrupt and opportunistic, where it avoids conspecifics and therefore hunts solitarily. This hunting behaviour is similar to other observed varanids (Mayes et al. 2005). We did not observe any encounter between varanid conspecifics or competition on prey-catching during the period of our work. This may be attributed to the social behaviour of varanids where they avoid encounters and conflicts with conspecifics (e.g., in Tsellarius and Tsellarius 1997). Most Varanus species prey on smaller vertebrates such as frogs and birds, but larger species can eat large animals as big as deer (Losos and Greene 1988). There are archaeological evidence shows large monitor lizards dwelling in caves (Price et al. 2015) but contemporary evidence on varanid feeding on caves bats has not previously been documented.

At present, we offer two plausible explanations to explain these hunting behaviour. First, we hypothesise that the recent influx of crows in the cave site may probably be explained by the ‘doomed surplus paradigm’, which predators take advantage and hunt the overcrowded population (Errington 1946). The assumption in this concept is the preyed upon individuals were low-quality populations, and the predators play a role to assure check and balance in the health of the population. Our observation on *V. cumingi* solely predates vulnerable individuals (i.e., weak or presumably sick, old, or injured) roosting on the lower part of the cave while *C. macrorhynchos* hunts on individuals that roost in abundant clusters. Secondly, in support of the first hypothesis, we suspect the population ‘spillover effects’ may have probably influenced the increase in bat biomass and predator influx in the cave. Although this concept is widely discussed and applied in marine reserves where no-take reserves increase and enhance the community complexity (e.g., species diversity and functional groups) due to the improvement of habitat quality and abundance of the prey availability and recruit predators (Russ and Alcala 2011). In the case of Monfort bat cave, predator recruitment is driven by the stable population of *R. amplexicaudatus* population due to absent of disturbances such as hunting and cave intrusion compared to most cave systems that experience high levels of disturbances (Tanalgo and Hughes, 2019). Here, we suggest that predators in Monfort Bat Cave are likely important to help maintain the ecological balance of the large aggregation of *R. amplexicaudatus* in the cave system. In addition to our description of predation by *C. macrorhynchos* and *V. cumingi*, pythons and large numbers of rats also live in the cave and have been observed to prey upon the bats. Moreover, we speculate another hypothesis that the presence of these predators may explain and shed light on the ‘anecdotally abnormal’ behaviour of *R. amplexicaudatus* in the cave in addition to overcrowding, where they remain aggressive most of the time during day time.

While our findings are fascinating, our work remains preliminary and we believe there are more that remains to be explored ahead. The predation of two predators on a cave-dwelling bat *R. amplexicaudatus* was described on this paper and offers another understanding of ecological and behavioural interactions present within caves and underground habitats. In addition, we have also shown here that modes of predation influence animal behaviour (e.g., the frequency of hunting and food intake) and information on the prey-predator dynamics on cave-roosting *R. amplexicaudatus* and offered a potential explanation on why the species behave and interact with its biotic environment.

## 4. Acknowledgments

This work is supported by the National Natural Science Foundation of China (Grant #: U1602265, Mapping Karst Biodiversity in Yunnan), Strategic Priority Research Program of the Chinese Academy of Sciences (Grant #: XDA20050202), Chinese Academy of Sciences Southeast Asia Biodiversity Research Center fund (Grant #: Y4ZK111B01), and SEABA: Southeast Asian Atlas of Biodiversity (Grant #: Y5ZK121B01). This is a part of the PhD project of KCT supported by the Xishuangbanna Tropical Botanical Garden, University of Chinese Academy of Sciences, and the Chinese Government Scholarship council, P.R. China. The authors are also grateful to Emerson Y. Sy (Philippine Center for Terrestrial and Aquatic Research) for verifying the varanid species and Monfort Bat Cave staff for their assistance during the observations and fieldwork in Monfort cave.

## References

Bednarz JC (1988) Cooperative hunting Harris’ hawks (*Parabuteo unicinctus*). Science 239: 1525–1527.

Boesch C (2002) Cooperative hunting roles among Tai chimpanzees. Human Nature, 13 : (1), 27–46.

Bowman R (2003) Apparent cooperative hunting in Florida scrub jays. The Wilson Bulletin 115: 197–199.

Carpenter E, Gomez R, Waldien DL, Sherwin RE (2014) Photographic estimation of roosting density of Geoffroys Rousette Fruit Bat *Rousettus amplexicaudatus* (Chiroptera: Pteropodidae) at Monfort Bat Cave, Philippines. Journal of Threatened Taxa 6: 5838–5844.

Csorba G, Rosell-Ambal G, Ingle N (2008) *Rousettus amplexicaudatus*. The IUCN Red List of Threatened Species 2008: e.T19754A9010480. http://dx.doi.org/10.2305/IUCN.UK.2008.RLTS.T19754A9010480.en. Downloaded on 18 January 2019.

DeVault TL, Rhodes Jr OE, Shivik JA (2003) Scavenging by vertebrates: behavioral, ecological, and evolutionary perspectives on an important energy transfer pathway in terrestrial ecosystems. Oikos 102: 225–234.

Dinets V (2015) Apparent coordination and collaboration in cooperatively hunting crocodilians. Ethology, Ecology and Evolution 27: 244–250.

Dinets V (2017) Coordinated hunting by Cuban boas. Animal Behavior and Cognition 4: 24–29.

Emery NJ, Clayton NS (2004) The mentality of crows: convergent evolution of intelligence in corvids and apes. Science 306 : 1903–1907.

Errington PL (1946) Predation and vertebrate populations. The Quarterly Review of Biology 21: 144–177.

Flaspohler DJ (2017) American Crows (*Corvus brachyrhynchos*) Enter Abandoned Mine to Hunt Bats. The Wilson Journal of Ornithology 129: 394–397.

Gese EM, Knowlton FF (2001) The role of predation in wildlife population dynamics. In: Ginnetr TF and SE Henke (eds) The Role of Predator Control as a Tool in Game Management. Texas Agricultural Research and Extension Center, Texas, pp 7–25.

Glen AS, Dickman R (2014) Carnivores of Australia: Past, Present and Future. CSIRO Publishing, Australia.

Goodwin D (1982) Crows of the world 2nd edition. British Museum Natural History, London.

Heaney LR, Dolar ML, Balete DS, Esselstyn JA, Rickart AE, Sedlock JL (2010) Synopsis of Philippine Mammals. The Field Museum of Natural History in cooperation with the Philippine Department of Environment and Natural Resources - Protected Areas and Wildlife Bureau. http://archive.fieldmuseum.org/philippine_mammals accessed December 10 2016.

Hernández DL, Mell JJ, Eaton MD (2007) Aerial predation of a bat by an American Crow. The Wilson Journal of Ornithology 119: 763–764.

Hiltunen T, Laakso J (2013) The relative importance of competition and predation in environment characterized by resource pulses–an experimental test with a microbial community. BMC Ecology 13: 29.

Koch A, Auliya M, Schmitz A, Kuch U, Böhme W (2007) Morphological studies on the systematics of South East Asian water monitors (*Varanus salvator* Complex): nominotypic populations and taxonomic overview. Mertensiella 16: e80.

Koch A, Auliya M, Ziegler T (2010) Updated checklist of the living monitor lizards of the world (Squamata: Varanidae). Bonn Zoological Bulletin 57: 127–136.

Kurosawa R, Kono R, Kondo T, Kanai Y (2003) Diet of jungle crows in an urban landscape. Global Environmental Research 7: 193–198.

Law SJ, De Kort SR, Bennett D, Van Weerd M (2018) Diet and Habitat Requirements of the Philippine Endemic Frugivorous Monitor Lizard *Varanus bitatawa*. Biawak 12: 12–22.

Laycock P (1982) Avian predation on cave dwelling insectivorous bats. Bokmakierie 34: 17–18.

Lee YF, Kuo YM (2001) Predation on Mexican free-tailed bats by peregrine falcons and red-tailed hawks. Journal of Raptor Research 35: 115–123.

Lefevre KL (2005) Predation of a bat by American Crows, *Corvus brachyrhynchos*. Canadian Field Naturalist 119: 443–444.

Lima SL, O’keefe JM (2013) Do predators influence the behaviour of bats? Biological Reviews 88: 626–644.

Losos JB, Greene HW (1988) Ecological and evolutionary implications of diet in monitor lizards. Biological Journal of the Linnean Society 35 : 379–407.

Marzluff JM, Angell T (2007) In the company of crows and ravens. Yale University Press.

Mayes PJ, Thompson GG, Withers PC (2005) Diet and foraging behaviour of the semi-aquatic *Varanus mertensi* (Reptilia: Varanidae). Wildlife Research 32: 67–74.

McFarland D (1981) The Oxford Companion to Animal Behavior. Oxford University Press, New York.

Mech LD, Boitani L (2003) Wolves: Behavior, Ecology, and Conservation. University of Chicago Press, Chicago.

Mikula P, Morelli F, Lučan RK, Jones DN, Tryjanowski P (2016) Bats as Prey of Diurnal Birds: A global perspective. Mammal Review 46: 160–174.

Packer, C. and L. Ruttan. 1988. The evolution of cooperative hunting. The American Naturalist 132: 159–198.

Price GJ, Louys J, Cramb J, Feng YX, Zhao JX, Hocknull SA, Joannes-Boyau R (2015) Temporal overlap of humans and giant lizards (Varanidae; Squamata) in Pleistocene Australia. Quaternary Science Reviews 125: 98–105.

Richardson H, Verbeek NA (1987) Diet selection by yearling northwestern crows (*Corvus caurinus*) feeding on littleneck clams (*Venerupis japonica*). The Auk 263–269.

Ritchie EG, Elmhagen B, Glen AS, Letnic M, Ludwig G, McDonald RA (2012) Ecosystem restoration with teeth: what role for predators? Trends in Ecology and Evolution 27: 265–271.

Roberts TW (1942) Behavior of Organisms. Ecological Monographs 12 : 339–412.

Russ GR, Alcala AC (2011) Enhanced biodiversity beyond marine reserve boundaries: The cup spillith over. Ecological Applications 21: 241–250.

Stimpson CM (2009) Raptor and owl bone from Niah Caves, Sarawak: identifications and morphological variation in the humerus and tarsometatarsus of selected raptors. International Journal of Osteoarchaeology 19: 476–490.

Sy E, Diesmos A, Jakosalem, PG, Gonzalez JC, Paguntalan LM, Demegillo A, Custodio C, Delima E, Tampos G, Gaulke M, Jose R (2009) Varanus cumingi. The IUCN Red List of Threatened Species 2009: e.T169897A6687602. http://dx.doi.org/10.2305/IUCN.UK.2009-2.RLTS.T169897A6687602.en. Downloaded on 19 January 2019.

Talbot LM (1978) The Role of Predators in Ecosystem Management. In: Holdgate MW, Woodman MJ (eds) The Breakdown and Restoration of Ecosystems. NATO Conference Series (Series I: Ecology), Vol 3. Springer, Boston, MA.

Tanalgo KC, Hughes AC (2018) Bats of the Philippine Islands – a review of research directions and relevance to national-level priorities and targets. Mammalian Biology 91: 46–56.

Tanalgo KC, Pineda JAF, Agravante ME, Amerol ZM (2015) Bird diversity and structure in different land-use types in lowland south-central Mindanao, Philippines. Tropical Life Sciences Research 26: 85–103.

Tanalgo KC, Hughes AC (2019) Priority-setting for Philippine bats using practical approach to guide effective species conservation and policy-making in the Anthropocene. Hystrix, the Italian Journal of Mammalogy

Tsellarius AY, Tsellarius EY (1997) Behavior of *Varanus griseus* during encounters with conspecifics. Asiatic Herpetological Research 7: 108–130.

Van Weerd M. et al. (2003) Flying foxes of the Northern Sierra Madre. *In* The Sierra Madre Natural Park, Northeast Luzon. In: Van der Ploeg, J., E.C. Bernardo and A.B. Masipiquena (eds.) Mountain range: global relevance, local realities. CVPED and Golden Press, Tuguegarao, the Philippines, pp. 54–62

Yosef R, Yosef N (2010) Cooperative hunting in brown-necked raven (*Corvus rufficollis*) on Egyptian mastigure (*Uromastyx aegyptius*). Journal of Ethology 28: 385–388.

